# Cathepsin B abundance, activity and microglial localisation in Alzheimer’s disease – Down syndrome and early onset Alzheimer’s disease; the role of elevated cystatin B

**DOI:** 10.1101/2023.06.20.545700

**Authors:** Yixing Wu, Paige Mumford, Suzanna Noy, Karen Cleverley, Alicja Mrzyglod, Dinghao Luo, Floris van Dalen, Martijn Verdoes, Elizabeth M.C. Fisher, Frances K. Wiseman

## Abstract

Cathepsin B is a cysteine protease that is implicated in multiple aspects of Alzheimer’s disease pathogenesis. The endogenous inhibitor of this enzyme, cystatin B (*CSTB)* is encoded on chromosome 21. Thus, individuals who have Down syndrome, a genetic condition caused by having an additional copy of chromosome 21, have an extra copy of an endogenous inhibitor of the enzyme. Individuals who have Down syndrome are also at significantly increased risk of developing early-onset Alzheimer’s disease (EOAD). The impact of the additional copy of cystatin B (*CSTB)* on Alzheimer’s disease development in people who have Down syndrome is not well understood. Here we compared the biology of cathepsin B and cystatin B (CSTB) in individuals who had Down syndrome and Alzheimer’s disease, with disomic individuals who had Alzheimer’s disease or were ageing healthily. We find that the activity of cathepsin B enzyme is decreased in the brain of people who had Down syndrome and Alzheimer’s disease compared with disomic individuals who had Alzheimer’s disease. This change occurs independently of an alteration in the abundance of the mature enzyme or the number of cathepsin B^+^ cells. We find that the abundance of cystatin B (CSTB) is significantly increased in the brains of individuals who have Down syndrome and Alzheimer’s disease compared to disomic individuals both with and without Alzheimer’s disease and we go on to investigate how this impacts enzyme activity in mouse and human cellular preclinical models of Down syndrome.

## Introduction

People who have Down syndrome (DS) are at much greater risk of developing early-onset Alzheimer’s disease (EOAD) than individuals from the general population, by the age of 50 around half of all people with DS will have been diagnosed with clinical dementia [1]. DS is caused by an extra copy of human chromosome 21 (Hsa21), which encodes ∼221 protein-coding genes (Ensembl release 109 - Feb 2023). The gene for Amyloid precursor protein (*APP*) is located on Hsa21, and duplication of this gene (*dupAPP*) is sufficient to cause EOAD in individuals who do not have DS [2] and is proposed to be both necessary and sufficient to trigger the early development of AD in individuals who have DS [3–5]. However, increasing evidence indicates that AD-relevant processes are modulated by Hsa21 trisomy; including the accumulation of amyloid-β [6–11] and the neuroinflammatory response to disease-associated pathology [12–16]. These differences are caused by the additional copy of gene(s) on the chromosome, resulting in elevated abundance of gene products that perturb molecular stoichiometry and lead to the changes in biology that occur in people who have DS. Understanding these differences in AD development in people who have DS is important for effective diagnosis and treatment, for this important group of individuals who are at greatly increased risk of dementia. Moreover, this research can provide unique mechanistic insight into the processes that contribute to disease pathogenesis.

Cathepsin B is a lysosomal cysteine protease, that has been implicated in important aspects of AD development, including the processing of APP, the degradation of amyloid-β and neuroinflammatory responses to disease involving lysosomal damage signals and the induction of the inflammasome pathway in microglia [17]. Raised levels of cathepsin B and increased activity are associated with AD and may be a potential therapeutic target, although some aspects of the role of the enzyme in disease are debated [18]. The abundance and activity of cathepsin B has been reported to be raised in DS fibroblasts but activity in the brain is not well understood [19]. It is notable that an endogenous inhibitor of cysteine cathepsins, cystatin B (*CSTB*) is encoded on Hsa21. How the additional copy of this gene influences the changes to cathepsin B that occur during AD pathogenesis in people who have DS is not well understood. Here we use a combination of studies in human post-mortem brain material and preclinical model systems to address this knowledge-gap.

## Materials and Methods

### Animal welfare and husbandry

All experiments were undertaken in accordance with the Animals (Scientific Procedures) Act 1986 (United Kingdom), after local institutional ethical review by the Medical Research Council, University College London and results are reported in accordance with ARRIVE 2.0 guidelines.

Mice were housed in individually ventilated cages (Tecniplast) with grade 5, autoclaved dust-free wood bedding, paper bedding and a translucent red “mouse house”. Free access to food and water was provided. The animal facility was maintained at a constant temperature of 19-23°C with 55 ± 10% humidity in a 12-hour light/dark cycle. Pups were weaned at 21 days and moved to standardised same-sex group housing with a maximum of 5 mice per cage.

The following mouse strains were used, here we show the abbreviated name and then the official name and unique Mouse Genome Informatics (MGI) identifier: *App^NL-F^* (App^tm2.1Tcs^, MGI:5637816), Dp1Tyb (Dp(16Lipi-Zbtb21)1TybEmcf, MGI5703798) Dp(10)2Yey (Dp(10Prmt2-Pdxk)2Yey, MGI:4461400) and Dp(17)3Yey (Dp(17Abcg1-Rrp1b)3Yey, MGI:4461398). All mouse strains were maintained by backcrossing males and females to C57BL/6J mice (imported from the Jackson Laboratory) and for all strains used backcrossing had taken place for more than ten generations. For Figure 6a-d Dp(10)2Yey mice were crossed with *App^NL-F/+^* animals over two-generations to generate all required genotypes from the second generation (wild-type, Dp(10)2Yey), *App^NL-F/NL-F^*, Dp(10)2Yey;*App^NL-F/NL-F^*). All other mouse cohorts (Fig. 6e-j) were generated by backcrossing to C57BL/6J mice. Animals were euthanized by exposure to rising carbon dioxide, followed by confirmation of death by dislocation of the neck in accordance with the Animals (Scientific Procedures) Act 1986 (United Kingdom).

### Culture of human fibroblasts

Human fibroblasts from individuals who had DS and age and sex-matched disomic controls (kindly supplied by the Coriell Biorepository) were grown in Dulbecco’s Modified Eagle Medium (DMEM) supplemented with 10 % fetal bovine serum and 1 % penicillin-streptomycin, at 37 °C with 5 % CO2. Cells were collected at 70 % confluency by trypsinisation (Gibco™ Trypsin-EDTA (0.25 %), Thermo Fisher Scientific) and mechanical agitation, pelleted, and washed 3 times in phosphate buffered saline (PBS) prior to homogenization. Experiments were undertaken between passage number 7 and 17. All experiments were blinded for trisomy of Hsa21 or disomic status.

### Cathepsin activity assays

Cathepsin B (CatB), and cathepsin D (CatD) endo-peptidase activity was determined using commercial assay kits (Cathepsin B: ab65300, Abcam; Cathepsin D: ab65302, Abcam). Samples were homogenized in Lysis buffer as per the manufacturer’s instructions, incubated on ice for 10 – 30 minutes, then centrifuged at 15000 *g* for 5 minutes. Protein concentration of the resultant supernatant was determined by Bradford assay. For CatB assays, 40-100 μg homogenate was diluted in 50 μl CB lysis buffer and loaded into 96 well microplates (Greiner Bio-One flat black clear bottom). For CatD, 5-10 μg homogenate was loaded. 2 μl CatB inhibitors N-Acetyl-L-leucyl-L-leucyl-L-methioninal (ALLM) (Abcam, ab141446) or Z-Phe-Phe-FMK (Abcam, ab141386), and 3 μg Pepstatin A (Sigma) (an inhibitor of aspartyl proteases including CatD) were used for negative controls. The substrate (Ac-RR) labelled with amino-4-trifluoromethyl coumarin (AFC) was used for the CatB assay; and for CatD, GKPILFFRLK(Dnp)-D-R-NH2 labelled with MCA was used. Plates were incubated at 37 °C in the dark and fluorescence activity measured at Ex/Em λ 400/505 nm (CatB) and Ex/Em λ 328/460 nm (CatD) every 90 seconds over a period of 135 minutes on a Tecan fluorometric plate reader.

The rate of reaction, proportional to enzymatic activity, was calculated using the slope of the linear phase of the reaction for a minimum of 10 cycles and expressed as a % of the equivalent slope for the control samples. For fibroblast samples and samples of human temporal cortex, an inhibited control was run for each individual sample used. For samples of mouse cortex, a representative inhibited control for each genotype was used. To control for non-specific activity, any activity detected in the appropriate inhibited control reaction was subtracted from the total activity observed in the uninhibited samples prior to analysis. Means of three technical replicates were calculated for each individual sample, with biological replicate being used as the experimental unit. For CatB assays 4, 10 and 2-6 technical replicates were used for human post-mortem temporal cortex, fibroblasts and mouse cortex samples respectively. For CatD assays 2 technical replicates were run for each human post-mortem temporal cortex sample.

### BMV109 staining and in-gel assay

For BMV109 staining of human fibroblasts, cells were incubated in Cellvis 6 Well glass bottom plates with high performance #1.5 cover glass (Fisher Scientific). For control wells to be pre-treated with cathepsin inhibitors, CA074ME (Merck Millipore, 10uM) was added 30 minutes prior to the BMV109 preincubation. BMV109 (A kind gift from Dr Martijn Verdoes, RIMLS, Netherlands, 1μM, 1:5,000) and LysoTracker™ Green DND-26 (Thermo Fisher Scientific, 50nM, 1:20,000) were then added for 1 hour and followed by 3 washes in PBS. Live cell imaging was then conducted by ZEISS LSM 980 confocal microscope (Carl Zeiss, Germany).

After confocal imaging, fluorescent SDS-polyacrylamide gel electrophoresis (SDS-PAGE) was conducted by following Edgington-Mitchell et al., (2016) [20] with minor modifications. In brief, fibroblasts were lysed in CB Cell Lysis Buffer from Cathepsin B Activity Assay Kit (Abcam, ab65300). NuPAGE LDS Sample Buffer (Thermo Fisher Scientific) and NuPAGE Sample Reducing Agent were added to the samples, which were then heated for 5 minutes at 95 °C. SDS-PAGE was conducted by running samples on NuPAGE Novex 4-12 % Bis-Tris Protein Gels in MOPs or MES SDS running buffer (both Thermo Fisher Scientific) at 120 V for 1.5 hours. After SDS-PAGE, the gel was taken to be imaged by Odyssey DLx Fluorescence Imaging System (LI-COR Biosciences). After imaging, the gel was transferred directly into the Bio-Safe™ Coomassie Stain (BIO-RAD, #1610786) and incubated overnight at 4 °C with agitation. The gel was then taken to be imaged by Odyssey DLx Fluorescence Imaging System (LI-COR Biosciences).

### Western blotting

Human fibroblasts were lysed in M-PER Mammalian Protein Extraction Reagent (Thermo Fisher Scientific) supplemented with cOmplete protease and PhosSTOP phosphatase inhibitors (Roche). Human temporal cortex samples were lysed in CB Cell Lysis Buffer from the Cathepsin B Activity Assay Kit (Abcam, ab65300). Mouse cortex samples were lysed in RIPA buffer (150 mm sodium chloride, 50 mm Tris, 1 % NP-40, 0.5 % sodium deoxycholate, 0.1 % SDS) plus complete protease inhibitors (Calbiochem) by mechanical disruption.

Samples were denatured in NuPAGE LDS Sample Buffer (Thermo Fisher Scientific) and NuPAGE Sample Reducing Agent in a 95 °C heat block for 5 minutes. Samples were then loaded into NuPAGE Novex 4-12 % Bis-Tris Protein Gels in MOPs or MES SDS running buffer (both Thermo Fisher Scientific) and SDS-PAGE was conducted by the application of 200 V for 50 minutes or 120 V for 1.5 hours. The proteins in the gel were transferred to a nitrocellulose membrane by Trans-Blot® Turbo™ Transfer System (Bio-Rad Laboratories) at 25 V, 2.5 A, for 7 minutes. The membranes were blocked with Intercept (PBS) Blocking Buffer (LI-COR Biosciences) for 1 hour at room temperature. The membranes were then incubated in primary antibodies in Intercept (PBS) Blocking Buffer (LI-COR Biosciences) overnight with agitation at 4 °C. The membranes were washed three times in PBST for 10 minutes before application of IRDye 680RD or 800CW secondary antibodies (LI-COR Biosciences) for 1 hour at room temperature: both at 1:10,000 dilutions in Intercept (PBS) Blocking Buffer (LI-COR Biosciences). After antibody probing, membranes were subjected to another three 10-minute washes with PBST, and images were then taken by Odyssey DLx Fluorescence Imaging System (LI-COR Biosciences). The density of protein bands was analysed with ImageJ. β-actin or GAPDH was used for the normalisation of CSTB and CatB protein levels. Relative intensities were calculated by dividing the lane density by the healthy ageing or WT within gel average density.

The primary antibodies are rabbit polyclonal anti-cystatin-B (Abcam, ab236646, 1:2,000), anti-cathepsin B (Abcam, ab92955, 1:1,000 and Merck, Ab-3 1:100) and anti-β-actin mouse monoclonal antibody (Sigma-Aldrich, #A5441, 1:5,000). The secondary antibodies were IRDye® 800CW Goat anti-Rabbit IgG (H + L) (1:10,000) and IRDye 680RD Goat anti-Mouse IgG (H+L) (1:10,000) (LI-COR Biosciences).

### Immunostaining (fibroblasts)

Human fibroblasts were cultured on coverslips in 12 or 24-well plates. Cells were first washed with PBS, then fixed with 4 % PFA for 10 minutes. After three PBS washes, cells were permeabilised with PBST containing 0.1 % Triton X-100 (Sigma-Aldrich). Cells were then blocked for 1 hour at room temperature in blocking solution containing 5 % goat serum, Tween-20 (Bio-Rad), and Glycine (Sigma-Aldrich) to prevent non-specific antibody binding. The cells were then stained with primary antibodies overnight at 4 °C in a humidified chamber. After primary antibody incubation, cells were washed with PBST, three times prior to one-hour secondary antibody incubation at room temperature. After secondary antibody incubation, cells were subjected to another three washes with PBST and mounted with Invitrogen ProLong Gold Antifade Mountant with DAPI (Thermo Fisher Scientific). Images were then taken by ZEISS LSM 980 confocal microscope (Carl Zeiss, Germany).

The primary antibodies are rabbit polyclonal anti-cystatin-B (Abcam, ab236646, 1:200), anti-cathepsin B (Abcam, ab92955, 1:500) and mouse anti-LAMP1 (Cell Signalling, #15665, 1:100). The secondary antibodies were Goat anti-Rabbit IgG (H+L) Highly Cross-Adsorbed Secondary Antibody, Alexa Fluor™ Plus 488 (Thermo Fisher Scientific, A32731, 1:700) and goat-anti-rabbit-IgG-H-L-highly-cross-adsorbed-secondary-antibody-polyclonal (Thermo Fisher Scientific, A11004, 1:700).

### Immunostaining (human post-mortem brain material)

To deparaffinise the samples, slides were first treated with xylene for three 5-minute intervals, followed by 20-30 dips in 100 %, 90 % and 70 % ethanol, and then 20-30 dips in water. Antigen retrieval was then carried out by placing the slides in 10mM citrate buffer, pH 6, and heated in a microwave for 5 minutes at high power. After cooling for 3 minutes, the slides were heated for 2 minutes at high power, followed by another 3 minutes at high power after cooling. The slides were then washed in water for 10 minutes with agitation, followed by 3 washes in PBST (PBS, 0.2 % triton X-100) for 10 minutes each with agitation. The samples were then preincubated with blocking buffer (PBS, 0.2 % triton X-100, and 5% Donkey serum) for 1 hour at room temperature in a humid chamber. Primary antibodies (goat anti-IBA1 (Abcam ab5076, 1:100) and rabbit anti-Cathepsin B (Calbiochem, #PC41, 1:100)) were added to the blocking solution and incubated overnight at 4 °C in the humid chamber. After overnight incubation, the samples were washed in PBS and 0.2 % triton for 3 times, 10 minutes each, and then secondary antibodies (Alexa Fluor 647 donkey anti-Rabbit (Invitrogen, #A31573, 1:500) and donkey anti-Goat IgG H&L Alexa Fluor 488 (Abcam, ab150129, 1:500)) diluted in blocking buffer were added to the samples and incubated for 1 hour at room temperature in the humid chamber. The samples were then washed in PBS and 0.2 % triton for 3 times, 10 minutes each, and DAPI was added for nuclear counterstain. The samples were incubated in DAPI solution for 5 minutes, then washed in PBS. Slides were incubated with TrueBlack Plus Lipofuscin Autofluorescence Quencher (Cambridge Bioscience, BT23014, 1:40 dilution in PBS) for 20 minutes, followed by washing in PBS. Then samples were mounted in ProLong Antifade Mountant (Thermo Fisher Scientific) and imaged by ZEISS LSM 980 confocal microscope (Carl Zeiss, Germany).

For each case two slides were stained and analysed. For each slide, 10 fields were chosen randomly and imaged using a 20 × objective lens with a 426.67 μm × 426.67 μm total size per image. The images were exported as TIF files. To count the cells, the Cell Counter plugin (FIJI for ImageJ) was used. Any cell with red or green pixels within a proximity of 20 µm to a DAPI stained nucleus was identified as a cathepsin B or IBA1 positive cell.

### Nuclear sub-fractionation

Nuclear and cytoplasmic sub-fractionations of human fibroblasts were carried out by using NE-PER™ Nuclear and Cytoplasmic Extraction Reagents (Thermo Fisher Scientific) as per the manufacturer’s instructions. In brief, human fibroblasts were resuspended by trypsinisation (Gibco™Trypsin-EDTA (0.25%) (Thermo Fisher Scientific)) and then centrifuged at 500 × g for 5 minutes. The cell pellet containing approximately 1×10^6^ cells was then washed with PBS prior to transferring to a 1.5mL Eppendorf tube. Cells were then centrifuged at 500 × g for 2-3 minutes. The supernatant was removed and the cell pellet resuspended in ice-cold CER I. The tube was then vortexed on the highest speed for 15 seconds and the resuspended cells incubated on ice for 10 minutes. Ice-cold CER II buffer was added and the tube vortexed again for 5 seconds. The tube was incubated on ice for 1 minute and then vortexed again for 5 seconds. The cell lysate was then centrifuged for 5 minutes at maximum speed (∼16,000 × g) at 4 °C. The supernatant, which contains the cytoplasmic fraction was transferred to a new pre-chilled tube and stored at -80 °C for further use. The remaining pellet fraction containing the nuclei was resuspended in ice-cold NER and vortexed again. The sample was placed on ice and the 15-second vortexing step was repeated every 10 minutes for 40 minutes in total. The tube was then centrifuged at maximum speed (∼16,000 × g) for 10 minutes at 4 °C. The supernatant that contained the nuclear extract fraction was transferred to a new pre-chilled tube.

Cytoplasmic and nuclear fractions were then subjected to western blotting and labelled with rabbit monoclonal anti-GAPDH (Sigma, G9545, 1:5,000) antibody and rabbit polyclonal anti-LSD1 antibody (Abcam, ab129195, 1:2,500).

### Experimental design and statistical analysis

Individual post-mortem brain donor, independent cell-line or individual animals were used as the unit of replication. Unique identification numbers were assigned to all samples to ensure the study remained blind and randomised during data acquisition and analysis. Statistical analysis was carried out using SPSS Statistics 22 (IBM) for ANOVA and Prism8 (GraphPad) for linear regression. When ANOVA was used for analysis, for studies of human post-mortem brain, gender and case type (healthy ageing, EOAD or AD-DS) were used as variables and age at death and post-mortem interval (PMI) as covariates; for mouse tissue studies genotype and sex of animal were used as variables. For fibroblast experiments, T-test was used for analysis with disomy/trisomy of Hsa21 status as the variable. The mean of technical replicates was used for biochemical datasets and cellular immunofluorescence data. Repeated measures ANOVA was used for immunofluorescence data from cases of human post-mortem brain, slide 1 and slide 2 treated as repeated measures. Graphs were plotted using Prism8 (GraphPad), data are presented as mean ± Standard Error of the Mean (SEM), p-values less than 0.05 are considered to be statistically significant.

## Results

### Case demographics

In this study we compared cases of EOAD (without known causal mutations in *APP, PSEN1* or *PSEN2*), Alzheimer’s disease in individuals who had DS (AD-DS), and healthy ageing disomic individuals. All cases were sourced from the South West Dementia Brain Bank (SWDBB), University of Bristol, UK and were matched as far as possible for demographics (Table 1, Supplementary table 1). The gender of individuals did not differ between case types (four male and six female cases per group). The cases of healthy ageing used in this study were significantly older than EOAD and AD-DS but no significant differences in age occurred between EOAD and AD-DS groups (ANOVA F(2,27) = 5.651, p = 0.009; pairwise comparison with Bonferroni correction healthy ageing compared with EOAD p = 0.028 with AD-DS p = 0.017). No significant difference in PMI was found between case type (ANOVA F (2,27) = 0.487, p = 0.620). No significant difference was observed in the number of cases that had a *APOE* 2/3, 2/4 alleles compared with 3/3 alleles or with 3/4, 4/4 alleles (*X^2^* (2, 29) = 1.945 p = 0.745). However, *APOE* 4/4 homozygotes were only observed in the EOAD group, likely contributing to the early development of disease in these individuals. The majority of cases of EOAD and AD-DS were Braak neurofibrillary tangle stage VI [21], one case of EOAD and two cases of AD-DS were Braak stage V, one case of AD-DS was Braak stage IV and insufficient material was available to stage one AD-DS case. In this study all samples were of temporal cortex Brodmann Area 21.

**Table 1.**
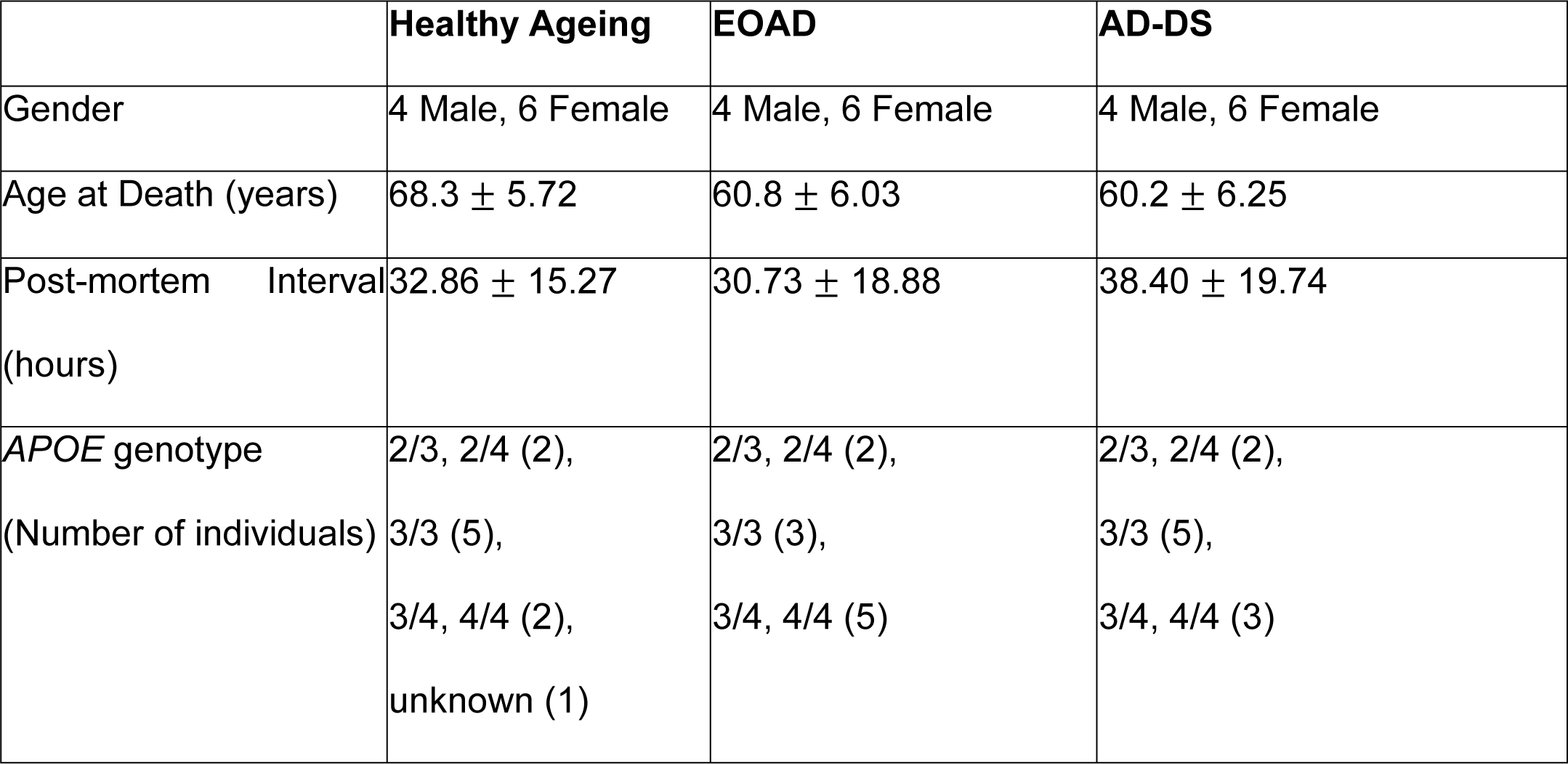

|  | Healthy Ageing | EOAD | AD-DS |
| --- | --- | --- | --- |
| Gender | 4 Male, 6 Female | 4 Male, 6 Female | 4 Male, 6 Female |
| Age at Death (years) | 68.3 ± 5.72 | 60.8 ± 6.03 | 60.2 ± 6.25 |
| Post-mortem Interval<br>(hours) | 32.86 ± 15.27 | 30.73 ± 18.88 | 38.40 ± 19.74 |
| *APOE* genotype<br>(Number of individuals) | 2/3, 2/4 (2),<br>3/3 (5),<br>3/4, 4/4 (2),<br>unknown (1) | 2/3, 2/4 (2),<br>3/3 (3),<br>3/4, 4/4 (5) | 2/3, 2/4 (2),<br>3/3 (5),<br>3/4, 4/4 (3) |

### Down syndrome Alzheimer’s disease (AD-DS) and matched disomic Alzheimer’s disease (EOAD) and healthy ageing case details

Summary of case demographics information of human post-mortem cases from matched disomic healthy ageing, EOAD and AD-DS cases, including age at death, post-mortem-interval and *APOE* genotype.

### Three copies of *CSTB* increase the abundance of CSTB protein in the brains of individuals with AD-DS compared with those who have EOAD

Hsa21 encodes the endogenous inhibitor of cysteine cathepsins, cystatin B gene (*CSTB)*. To determine if trisomy 21 results in an increase in cystatin B protein (CSTB) in the brains of people who have AD-DS, we compared the abundance of the protein between these individuals and individuals from the general population with EOAD or who were ageing healthily, by western blot. The abundance of cystatin B (CSTB) was significantly higher in cases of people who had AD-DS than in those from the general population who had EOAD or those that did not have dementia (healthy ageing) (Fig. 1a-d). The age and sex of the individual and the post-mortem interval did not significantly affect cystatin B (CSTB) abundance. Thus, three-copies of *CSTB* resulting from trisomy of Hsa21, is associated with increased cystatin B (CSTB) protein in the brains of people who have AD-DS compared to disomic individuals who have EOAD or are ageing healthily.

**Figure 1.**
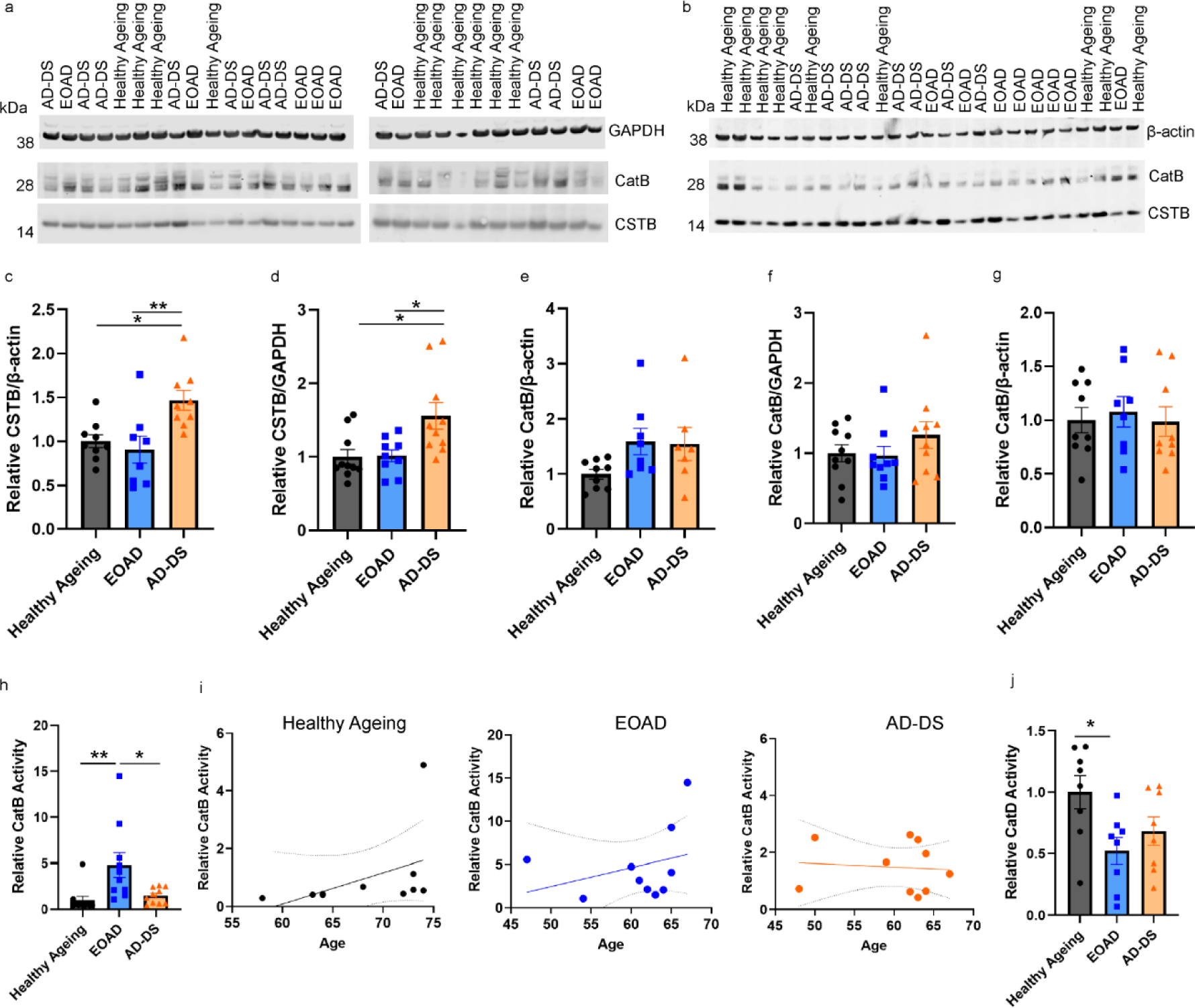
Cystatin B (CSTB) abundance is increased and cathepsin B activity is reduced in the brains of individuals with AD-DS, independently of changed cathepsin B protein level or cathepsin D activity. Cystatin B (CSTB) (a-d) and cathepsin B (a, e, f Ab-3 or b, g ab92955), protein abundance measured by western blot in human temporal cortex from people who had AD-DS, EOAD and healthy ageing. (c, d) Type of case altered the abundance of cystatin B (CSTB) protein (ANOVA β-actin F (2,18) = 9.087, p = 0.002; GAPDH F (2,21) = 7.500, p = 0.003). Cystatin B (CSTB) abundance was higher in individuals who had AD-DS than those with EOAD (pairwise comparisons with Bonferroni correction β-actin p = 0.008, GAPDH p = 0.020) and healthy controls from the general population (pairwise comparisons with Bonferroni correction, β-actin p = 0.024; GAPDH p = 0.013), with no difference in abundance between individuals with EOAD and control individuals (pairwise comparisons β-actin and GAPDH p = 1.000). (e-g) The abundance of mature cathepsin B protein, (e, f Ab-3 or g ab92955), was not altered by the type of case (Ab-3 ANOVA β-actin F (2,16) = 3.051, p = 0.075; GAPDH F (2,21) = 2.141, p = 0.143) (ab92955 ANOVA F (2,18) = 0.970, p = 0.398). (h) Type of case affected cathepsin B activity (ANOVA F(2,22) = 9.027, p < 0.001); activity was significantly higher in individuals who had EOAD than in controls (pairwise comparisons with Bonferroni correction p = 0.004), but lower in individuals with AD-DS than those with EOAD (pairwise comparisons with Bonferroni correction, p = 0.012), with no difference between individuals with AD-DS and control individuals (pairwise comparisons p = 1.000). The age (in years) at time of death significantly affected activity (ANOVA F (1,22) = 4.642, p = 0.042), (i) Within case types, no significant correlations were observed (Healthy ageing p = 0.199, EOAD p= 0.376 and AD-DS p = 0.807); PMI did not significantly alter activity (ANOVA F (1,22) = 0.946, p = 0.341). (j) Type of case significantly affected cathepsin D activity (ANOVA F(2,16) = 6.360, p = 0.009); activity was significantly lower in individuals who had EOAD than in healthy controls (pairwise comparisons with Bonferroni correction p = 0.022), no significant difference in activity was observed between AD-DS and healthy ageing controls (pairwise comparisons with Bonferroni correction, p = 0.180), with no difference between in individuals with AD-DS those with EOAD (pairwise comparisons p = 1.000). Individual data points are technical means for independent biological samples, error bars SEM. * p < 0.05 and ** p < 0.01.

### Down syndrome does not alter the abundance of mature cathepsin B in the brain but cathepsin B activity is lower in AD-DS than EOAD

In the same post-mortem samples, we quantified the abundance of mature cathepsin B protein (∼28 kDa), by western blot using two independent antibodies (Ab-3 and ab92955). We found no significant change in the abundance of mature cathepsin B protein between the subgroups of cases with either antibody (Fig. 1a, b, e, f). Post-mortem interval, age and gender did not significantly affect the abundance of mature cathepsin B.

As cystatin B (CSTB) protein abundance was elevated in individuals who had AD-DS we went on to investigate if cathepsin B endo-peptidase activity was impacted by the increase in the endogenous inhibitor. Using the same samples, we measured cathepsin B activity by biochemical assay; quantifying the rate of cleavage of Ac-RR labelled with amino-4-trifluoromethyl coumarin (AFC). To control for the non-specific activity, we measured the rate of cleavage in each sample inhibited by FMK(z-FA-FMK), a cysteine protease inhibitor. This non-specific activity was then subtracted from the total activity to identify the proportion of the activity specific to cathepsin B. Using this method, we observed that cathepsin B activity is elevated in the temporal cortex of people who had EOAD compared with healthy controls, but that no difference in cathepsin B activity was observed in individuals who have AD-DS compared to healthy ageing controls (Fig. 1h). Moreover, significantly less cathepsin B activity was measured in the temporal cortex of individuals who have AD-DS compared with individuals from the general population who had EOAD. Notably, consistent with other reports [22], we observed that the age of an individual significantly affects cathepsin B activity. However, the weak positive correlation between activity and age occurring in healthy ageing and EOAD subgroups was not statistically significant. Moreover, in contrast to the trend for a positive relationship between age and activity observed in the general population, we observed a non-significant inverse relationship between activity and age in individuals who had AD-DS (Fig. 1i).

To understand if the change in cathepsin B activity was specific, we assayed the activity of another major lysosomal protease (cathepsin D); the activity of which is not regulated by cystatin B (CSTB) [18]. We measured cathepsin D activity by the rate of cleavage of GKPILFFRLK(Dnp)-D-R-NH2 labelled with 7-methoxycoumarin-4-acetic acid (MCA). To control for non-specific activity, we measured the rate of cleavage in each sample inhibited by Pepstatin A, a cathepsin D inhibitor. This non-specific activity was then subtracted from the total activity to identify the proportion of the cleavage specific to cathepsin D. We found that cathepsin D activity was significantly decreased in the temporal cortex of individuals who had EOAD compared to the activity observed in healthy ageing control individuals (Fig. 1j). A similar pattern of decreased cathepsin D activity was also observed in samples of temporal cortex from individuals who have AD-DS, but this did not reach statistical significance. No difference in activity was detected between individuals who had AD-DS and those who had EOAD. These data are consistent with a decrease in cathepsin D activity reported in DS patient fibroblasts [19].

These data indicate that having three copies of *CSTB*, raises the abundance of cystatin B (CSTB) protein in the brains of individuals who have AD-DS. This may cause the specific decrease in cathepsin B activity observed in the brains of people who have AD-DS compared to the raised activity of the enzyme which occurs in people who have EOAD.

### Differences in cathepsin activity between EOAD and AD-DS are not the result of differences in Braak neurofibrillary tangle stage

In the case series studied, two cases of AD-DS were Braak neurofibrillary tangle stage V, one case was Braak stage IV and one case has not been staged. In comparison, only one EOAD case was Braak V and the other EOAD cases were Braak VI. To determine if differences in Braak tangle stage contributed to the difference in cathepsin activity between EOAD and AD-DS, we repeated our analysis comparing AD-DS (Braak VI, 6 cases) with EOAD (Braak VI, 9 cases) and healthy ageing (Braak 0-II, 10 cases). In this sub-analysis, cathepsin B activity was significantly different between case types (ANOVA F(2,17) = 6.792, p = 0.007), with activity being significantly higher in EOAD compared with healthy ageing (pairwise comparison with Bonferroni p = 0.008) but not different between AD-DS and healthy ageing (pairwise comparison with Bonferroni p = 1.000) (Supplementary Figure 1a). In this sub-set of the cases, cathepsin D activity significantly differed between case types (ANOVA F(2,14) = 7.188, p = 0.007), with activity being significantly lower in EOAD compared with healthy ageing (pairwise comparison with Bonferroni p = 0.028), a trend for reduced activity in AD-DS compared with healthy ageing (pairwise comparison with Bonferroni p = 0.070) but no difference in the activity being observed between AD-DS and EOAD (pairwise comparison with Bonferroni p = 1.000) (Supplementary Figure 1b). Thus, the difference in cathepsin B activity we observed in the cases of AD-DS compared with EOAD is likely to be caused by a specific effect of three copies of Hsa21 on enzyme function and not the result of differences in tau pathology in the cases.

### The number of IBA^+^ microglia and cathepsin B^+^ cells are upregulated in individuals who had EOAD and those who have AD-DS

Recent data has shown that the biology of microglia differs in people who have DS both prior to and after the development of AD pathology [13, 14, 23]. As cathepsin B is upregulated in microglia during the development of AD, we investigated if the changes in cathepsin B activity we observe in individuals who have DS, are related to changes in microglia. To identify microglia, we used an antibody against Ionized calcium binding adaptor molecule 1 (IBA1), which is expressed by microglia and macrophages. We quantified the number of IBA1^+^ microglia, the number of cathepsin B^+^ cells, the proportion of cathepsin B colocalised with IBA1^+^ microglia and the relative proportion of IBA^+^ microglia that contain cathepsin B, in samples of temporal cortex from people who had EOAD, AD-DS or healthy ageing (Fig. 2). We find that the number of IBA1^+^ cells is significantly increased in people who have EOAD and those with AD-DS compared with those undergoing healthy ageing (Fig 2d). However, no difference in the total number of IBA1^+^ cells was observed when people who had AD-DS and those who had EOAD were compared. Similarly, the number of cathepsin B^+^ cells is significantly increased in people who had EOAD or AD-DS compared with those who were undergoing healthy ageing but no difference in the number of cathepsin B positive cells occurred between individuals who had AD-DS and EOAD (Fig 2e).

**Figure 2.**
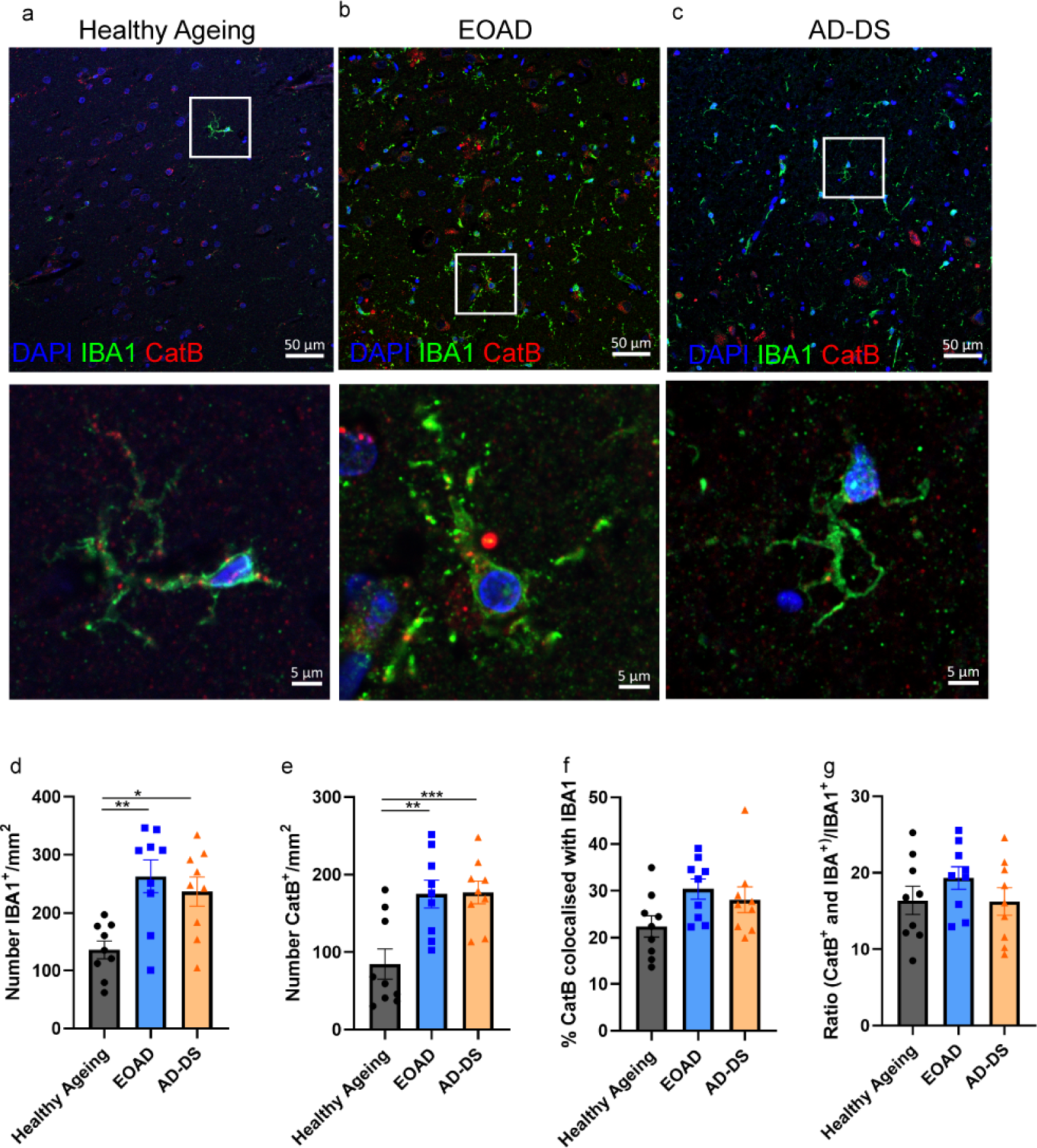
The number of IBA microglia and cathepsin B cells is increased in EOAD and AD-DS compared with healthy ageing but no change in the proportion of cathepsin B positive microglia occurs. (**a-c**) Representative images of IBA1 (green), cathepsin B (red) and DAPI-nucleus (blue) stained cells, and colocalised IBA1 and cathepsin B staining in human temporal cortex from individuals with (a) healthy ageing, (b) EOAD or (c) AD-DS. (**d**) The number of IBA1^+^ microglia/mm^2^ was significantly altered by the type of case (ANOVA F (2,19) = 6.527, p = 0.007); significantly more cells were observed in both cases of EOAD and AD-DS compared with healthy ageing (pairwise comparisons with Bonferroni correction EOAD p = 0.005, AD-DS p = 0.028) but no difference was observed between EOAD and AD-DS. (**e**) The number of cathepsin B^+^ cells/mm^2^ was significantly altered by the type of case (ANOVA F (2,19) = 13.379, p < 0.001); significantly more cells were observed in both cases of EOAD and AD-DS compared with healthy ageing controls (pairwise comparisons with Bonferroni correction EOAD p = 0.002, AD-DS p < 0.001) but no difference was observed between EOAD and AD-DS. (**f**) A trend to significance was observed between case types for the proportion of total cathepsin B staining that colocalised with IBA1^+^ microglia (ANOVA F (2,19) = 3.446, p = 0.053) but pairwise differences were not significant (pairwise comparisons with Bonferroni correction EOAD versus healthy p = 0.233, AD-DS versus healthy p = 0.398). (**g**) The proportion of IBA1 microglia that were also positive for cathepsin B was not altered by case type (ANOVA F(2,19) = 1.493, p = 0.250).

We observed a trend for case type to alter the proportion of total cathepsin B staining that colocalised with IBA1^+^ but the observed differences between the groups were not significant (Fig. 2f). Notably, the majority (∼70%) of cathepsin B staining did not colocalise with IBA1, indicating that the bulk of activity in the brain likely occurs in other cell-types (including microglia that do not express IBA1) or extracellularly. Moreover, case type did not affect the proportion of IBA^+^ microglia cells that were positive for cathepsin B (Fig. 2g). These data indicate that the differences observed in the activity of cathepsin B enzyme in people who have AD-DS are not related to the number of cells expressing the enzyme, consistent with our western blotting data. Nor does the distribution of the enzyme in microglia differ between individuals who have EOAD and AD-DS. Thus, the specific reduction in activity in individuals who have DS, likely occurs by an alternative mechanism and is not caused by a change in the distribution of the enzyme to microglia cells.

### Trisomy Hsa21-associated cathepsin B deficits are not modelled in human fibroblasts, despite the raised abundance and unchanged subcellular distribution of CSTB

To test the hypothesis that raised cystatin B/CSTB, caused by an additional copy of Hsa21 causes the changes to cathepsin B activity in the brains of individuals who have AD-DS, we measured CSTB and cathepsin B protein abundance, and cathepsin B activity in fibroblasts isolated from infants and children who have DS (trisomy 21) and matched disomic individuals (Supplementary table 2). We first determined if three copies of *CSTB* lead to the raised abundance of cystatin B (CSTB) protein, by western blot of total cellular proteins (Fig. 3a, b). Consistent with our results from human post-mortem brain material, an additional copy of Hsa21 significantly increased cystatin B (CSTB) protein in fibroblasts, isolated from individuals who have DS.

**Figure 3.**
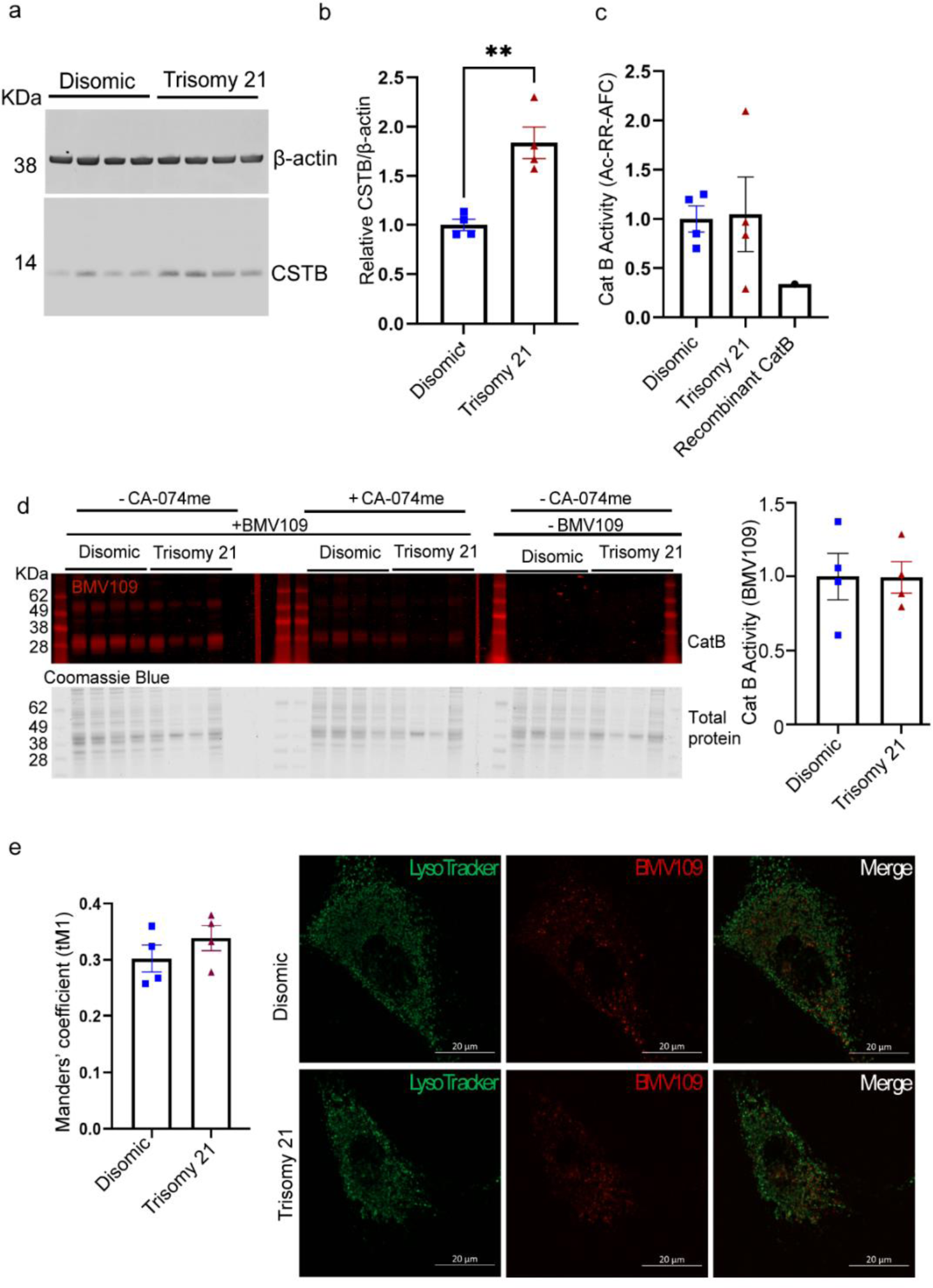
Increased abundance of cystatin B (CSTB) protein in fibroblasts isolated from individuals who have trisomy of Hsa21 is not sufficient to decrease total or lysosomal cathepsin activity compared to disomic matched cells. Cathepsin B activity and cystatin B (CSTB) protein abundance were measured in human primary fibroblasts (**a, b**) Trisomy of chromosome 21 significantly increases cystatin B (CSTB) protein abundance (T-test p = 0.003). (**c**) Trisomy of chromosome 21 does not alter total cathepsin B activity as measured by biochemical assay (rate of cleavage of Ac-RR-AFC), (T-test, p = 0.908) or (**d**) by BMV109 fluorescence (red) quantified by in gel assay based upon enzyme identification by molecular weight (28 kDa) normalized to total protein measured by Coomassie blue (T-test, p = 0.334). (**e**) Trisomy of Hsa21 does not alter cathepsin activity within lysosomes as measured by lysotracker (green) colocalised BMV109 fluorescence (red) (T-test, p = 0.310). Individual data points are group means for n = 4 disomic and n = 4 trisomy 21, independent cells lines (three technical replicates for western blots and 15-20 cells per lines for immunofluorescence), error bars SEM, ** p < 0.01.

Total cellular cathepsin B activity was measured using a biochemical fluorescence cleavage assay and no significant difference was observed between disomic and trisomy 21 cells (Fig. 3c). To investigate this further we assayed the activity of cathepsin B in disomic compared to trisomy 21 cells, using the activity-based probe BMV109 [24] (Fig. 3d). No significant difference in either total (Fig. 3d) or lysosomal (Fig. 3e) cathepsin B activity was observed between trisomy 21 and disomic fibroblasts using this method. This suggests that in fibroblast cells, three copies of Hsa21 raises cystatin B (CSTB) protein levels, but this is not sufficient to modulate cathepsin B activity.

Cystatin B (CSTB) has been reported to be localised to the cytosol, nucleus and lysosome [25, 26]; the pattern of subcellular localisation of the protein is critical to its regulation of cathepsin activity. Thus, we determined if three copies of Hsa21 altered the subcellular distribution of cystatin B (CSTB) and hence its impact on cathepsin activity, using biochemical subcellular fractionation and western blotting. We found no difference in the relative abundance of cystatin B (CSTB) in the nucleus and cytoplasm in trisomy 21 compared to disomic fibroblasts and confirmed that total abundance of the protein was raised by three copies of Hsa21 (Fig. 4a-c). We determined if trisomy 21 altered the relative amount of cystatin B (CSTB) that colocalised with LAMP1 (a lysosomal marker) in fibroblasts (Fig. 4d-e), as cathepsin activity in lysosomes is likely to have a particularly important role in APP processing and amyloid-β catabolism. We found no difference in colocalisation of cystatin B (CSTB) and LAMP1 between trisomy 21 and disomic cells.

**Figure 4.**
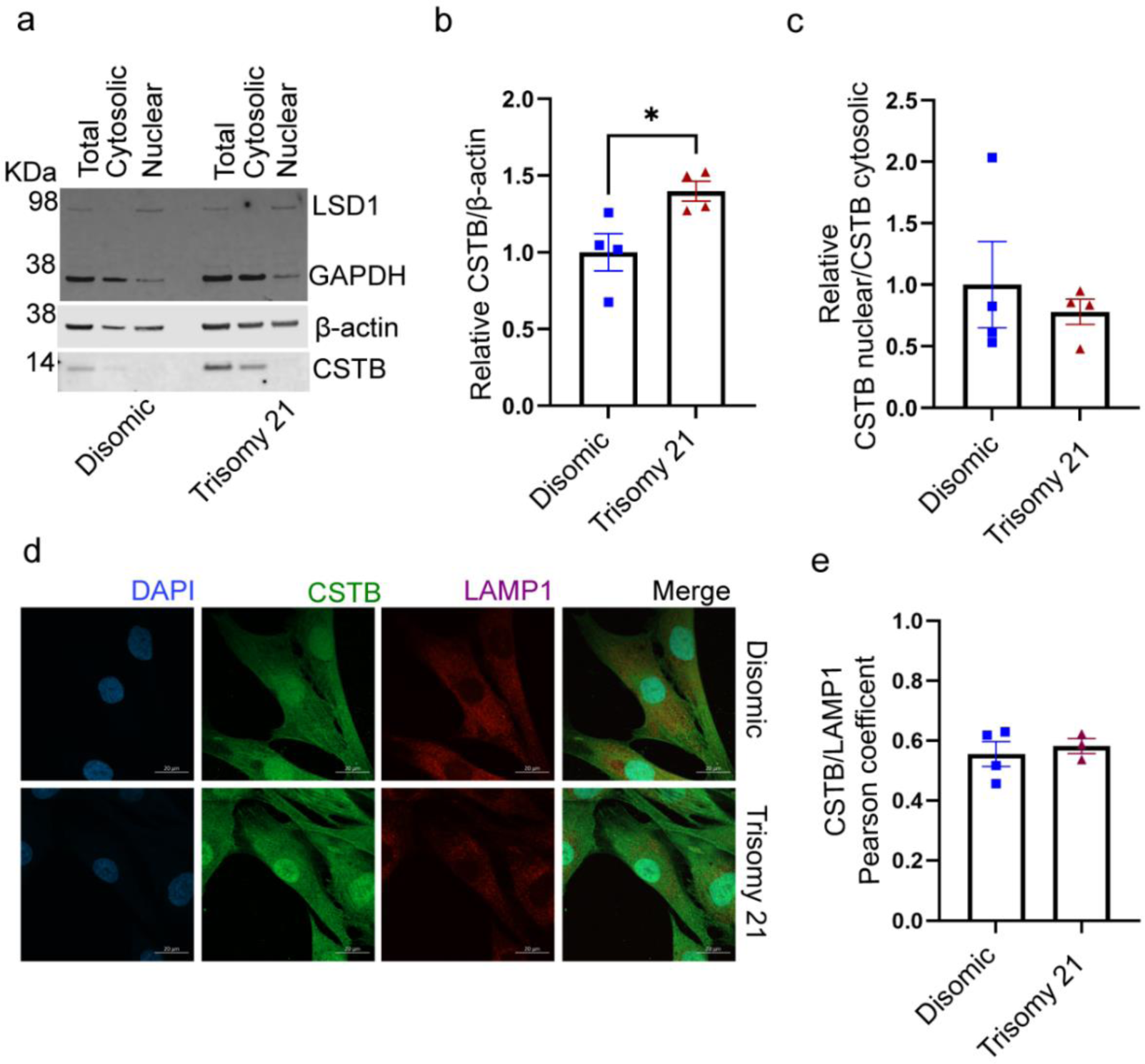
Trisomy of Hsa21 does not alter the localisation of cystatin B (CSTB) to the lysosome. **(a-c)** Total cellular proteins from disomic and trisomy 21 human primary fibroblasts were separated into cytosolic and nuclear fraction and the abundance of cystatin B (CSTB) quantified by western blot, LSD1 was used as a marker of the nuclear fraction. (**b**) Relative cystatin B (CSTB) abundance was increased by trisomy 21 (T-test p = 0.028) but the nuclear/cytosolic cystatin B (CSTB) ratio was not altered (T-test p = 0.567). (**d-e**) Colocalisation of cystatin B (CSTB) with LAMP1 did not differ between disomic and trisomy 21 cells (T-test p = 0.636). For d and e, individual data points are technical means of 3 technical replicates for n = 4 disomic and n = 3 trisomy 21, independent cell lines, error bars SEM. * p < 0.05.

To determine if raised cystatin B (CSTB) led to a compensatory alteration in cathepsin B protein abundance or processing; we quantified the amount of pro- and mature cathepsin B, generated by cleavage of the pro-form in trisomy 21 and disomic fibroblasts by western blotting. No differences in the relative abundance of pro- or mature cathepsin B were detected (Fig. 5a-c). Changes in the processing of cathepsin B were also not detected as measured by the ratio of the mature to pro-forms of the protein (Fig. 5d). We also determined if trisomy 21 altered the colocalization of cathepsin B with LAMP1 in fibroblasts (Fig. 5 e,f). We found no difference in the proportion of cathepsin B colocalised with LAMP1 between disomic and trisomy 21 cells (Fig. 5 e,f). Thus, the changes in cathepsin activity in the brains of people who have AD-DS are not modelled in a fibroblast model of trisomy of Hsa21 and may be specific to the central nervous system.

**Figure 5.**
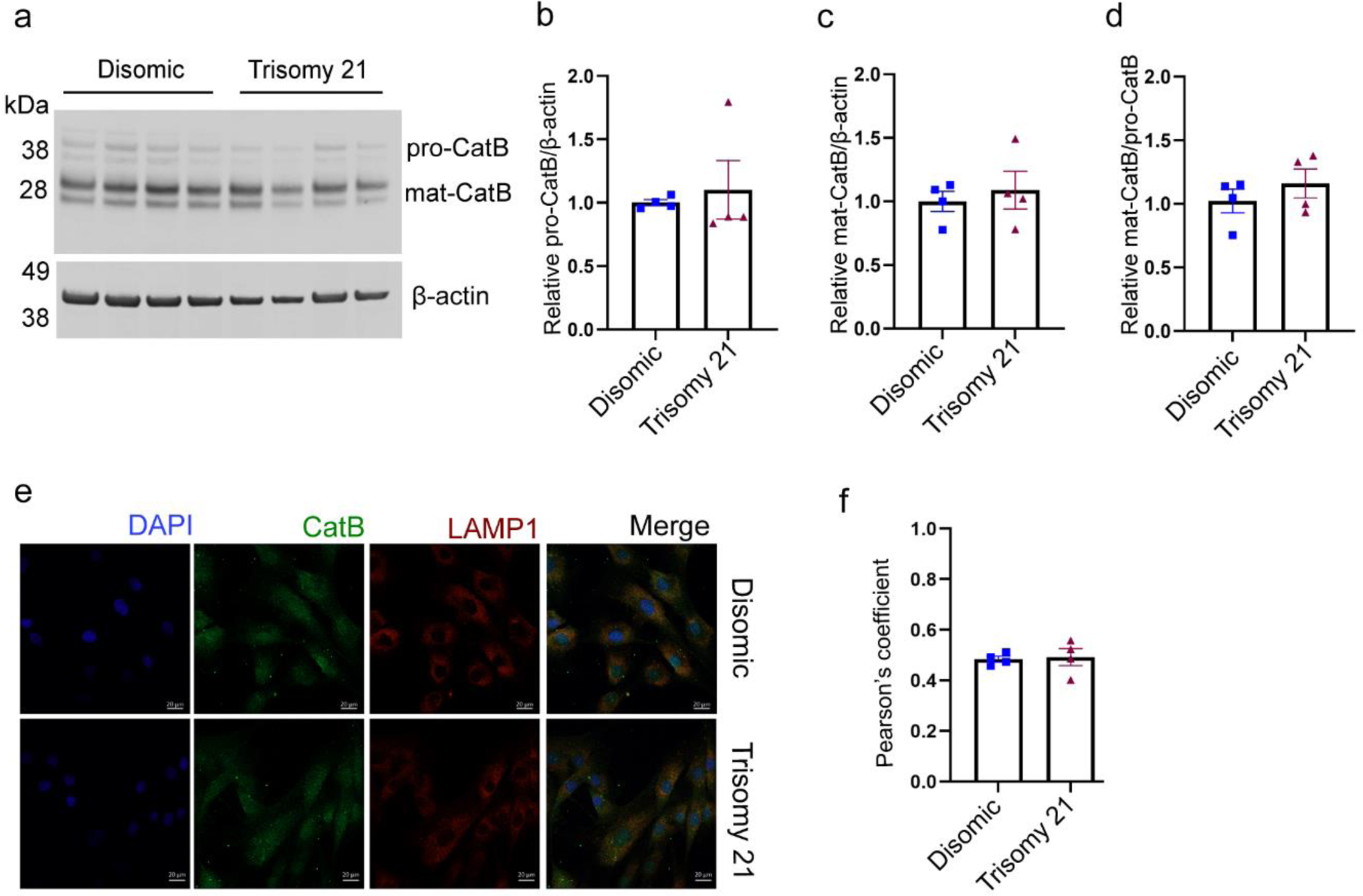
Trisomy of Hsa21 does not alter cathepsin B maturation or lysosomal localisation. (**a**) Western blot showing pro-cathepsin B (pro-CatB) and mature cathepsin B (mat-CatB) protein levels in disomic and trisomy 21 human primary fibroblasts (**b**) pro-cathepsin B protein levels (T-test p = 0.676) (**c**) mature cathepsin B protein level (T-test p = 0.613) (D) mature cathepsin B/pro-cathepsin B ratio (T-test p = 0.383) were not altered by trisomy 21. (**e-f**) Colocalisation of cathepsin B with LAMP1 was measured using immunofluorescence and no difference between disomic and trisomy 21 fibroblasts was observed (T-test p = 0.822). Individual data points are technical means of 3 technical replicates for n = 4 disomic and n = 4 trisomy 21 independent cell lines, error bars SEM.

### Three copies of *Cstb* increase CSTB abundance in the cortex of mouse models of AD-DS but do not alter Cathepsin B activity

To determine if an additional copy of *Cstb* is sufficient to raise cystatin B (CSTB) protein abundance or alter cathepsin B enzyme activity in the adult brain of a mouse model of DS, we measured CSTB protein level and cathepsin activity in the Dp(10)2Yey mouse model, which has an additional copy of 38 genes, carried on an internal duplication on Mmu10 of the region of homology with Hsa21. This region includes *Cstb* and thus the Dp(10)2Yey model carries three copies of this gene. We undertook the study in mice from a cross of this duplicated region with a mouse model (*APP^NL-F/NL-F^*) in which the endogenous mouse *App* gene is partially humanised and carries AD causal point mutations, resulting in the accumulation of human amyloid-β within the brain of the model (Supplementary table 3) [27]. We note that we have previously studied APP processing and amyloid-β aggregation and accumulation in this cross and found that these are not significantly altered [8].

Cystatin B (CSTB) protein level was significantly increased in the cortex of 3-month old Dp(10)2Yey mice; but was not altered in littermates by the expression of human APP/amyloid-β (Fig. 6a, b). We then went on to determine if endo-peptidase activity of cathepsin B was altered at 3-months of age in the cortex of a progeny from our cross of the DS and amyloid-β accumulation models, using the same methods as for our study of human-post-mortem brain. We found no consistent effect of the Dp(10)2Yey duplication on enzyme activity, although in one experiment a modest and significant decrease in activity was observed, but this result was not replicated (Fig. 6c-f).

**Figure 6.**
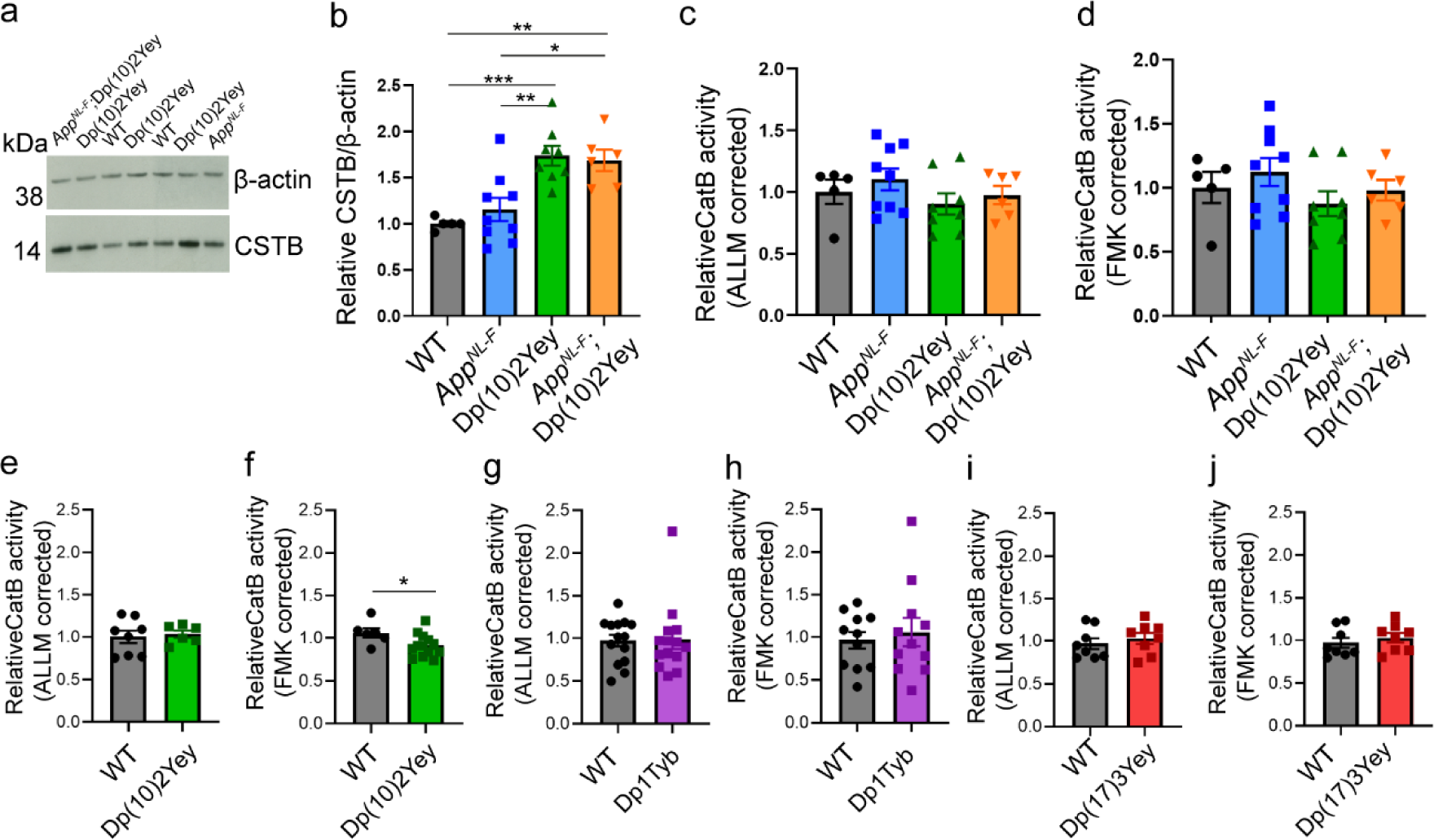
Three copies of *Cstb* increase cystatin B (CSTB) in the cortex of mouse models of AD-DS but do not alter cathepsin B activity. **(a, b)** Cystatin B (CSTB) protein abundance (determined by western blot) was measured in cortex of a 3-month old mouse model of AD-DS (progeny from a cross of Dp(10)2Yey and *App^NL-F/NL-F^*) which has three copies of *Cstb*. The abundance of cystatin B (CSTB) was significantly increased by the Dp(10)2Yey region (ANOVA F(1,22) = 31.269, p < 0.0001). Significantly higher cystatin B (CSTB) levels were detected in Dp(10)2Yey compared to wildtype (WT) (pairwise comparisons with Bonferroni correction, p < 0.001) and *App^NL-F/NL-F^* (pairwise comparisons with Bonferroni correction, p = 0.003) cortex. Significantly higher cystatin B (CSTB) levels were detected in Dp(10)2Yey; *App^NL-F/NL-F^* compared to wildtype (WT) (pairwise comparisons with Bonferroni correction, p = 0.005) and *App^NL-F/NL-F^* (pairwise comparisons with Bonferroni correction, p = 0.015) cortex. (**c-h**) Cathepsin B activity determined by biochemical assay (rate of cleavage of Ac-RR-AFC, corrected for non-specific activity measured in samples inhibited by (**c, e, g, i**) ALLM or (**d, f, h, j**) FMK(z-FA-FMK). (**c, d**) No difference in cathepsin B activity was detected in cortex of 3-month old mice with the Dp(10)2Yey segmental duplication compared to littermates with this duplication and the *App^NL-F/NL-F^* allele that results in amyloid-β accumulation (**c**) ALLM (ANOVA F(1,22) = 1.325 p = 0.262) or (**d**) FMK (ANOVA F(1,22) = 1.142 or p = 0.297). No difference in cathepsin B activity was detected in cortex from 3-month-old mice homozygous for the *App^NL-F^* allele (**c**) ALLM (ANOVA F(1,22) = 0.571 p = 0.458) or (**d**) FMK (ANOVA F(1,22) = 0.815 or p = 0.376). (**e**) No difference in cathepsin B activity was detected in cortex of 3-month old mice with the Dp(10)2Yey segmental duplication ALLM (ANOVA F(1,10) = p = 0.118). (**f**) A modest reduction in cathepsin B activity was detected in cortex of 3-month old mice with the Dp(10)2Yey segmental duplication FMK (ANOVA F(1,15) = 5.542 p = 0.033). No difference in cathepsin B activity was detected in cortex of 3-month-old mice with the Dp1Tyb segmental duplication (**g**) ALLM (ANOVA F(1,24) = 0.010 p = 0.923) or (**h**) FMK (ANOVA F(1,18) = 0.013 or p = 0.909). (**i, j**) No difference in cathepsin B activity was detected in cortex of 3-month old mice with the Dp(17)3Yey segmental duplication (**I**) ALLM (ANOVA F(1,12) = 0.543 p = 0.475) or (**J**) FMK (ANOVA F(1,12) = 0.731 or p = 0.409). Individual data points are technical means for independent biological samples, error bars SEM. * p < 0.05, ** p < 0.01, *** p < 0.001. Male and female mice were used and sex was included as a variable in the ANOVA, cohort details Supplementary table 3.

To determine if genes other than *CSTB* on Hsa21 when in three copies might be sufficient to modulate the activity of cathepsin B in mouse brain, we measured activity in the cortex of 3-month of age Dp1Tyb mouse model of DS, which has an additional copy of 148 Hsa21 homologous genes, including *App,* located on Mmu16 and the Dp(17)3Yey mouse model of DS, which has an additional copy of 17 Hsa21 homologues located on Mmu17. Enzyme activity was not altered by the presence of either segmental duplication, we note neither of these models carries an additional copy of *Cstb*. These data indicate that three copies of *Cstb* is not sufficient to modulate cathepsin B activity in a mouse model of AD-DS, consistent with our previous findings from a segmental duplication model of *Cstb* [28], nor is an additional copy of Hsa21 homologues encoded on Mmu16 or Mmu17 sufficient to alter activity.

## Discussion

Our data demonstrate that three copies of Hsa21 raises the abundance of cystatin B (CSTB) protein in the brains of individuals who have AD-DS, in the Dp2(10)Yey mouse model of DS, and in fibroblasts isolated from children who have DS compared with matched disomic controls. Increased abundance of this endogenous inhibitor of cysteine cathepsins could modulate key aspects of neurobiology by the inhibition of the activity of cathepsin B. This may be particularly relevant in AD pathogenesis in which abundance and activity of the enzyme is upregulated and has been suggested to impact on a number of key disease mechanisms, including APP processing, amyloid-β catabolism, and the neuroinflammatory response to pathology; particularly via the activation of inflammasomes [29–31].

Here, we demonstrate that the number of cathepsin B^+^ cells, and the activity of cathepsin B is significantly increased in people who have EOAD compared with those undergoing healthy ageing, supporting the proposed role of the enzyme as a potential therapeutic target in AD. Although the number of cathepsin B^+^ cells is similarly increased in individuals who have AD-DS, we find that the activity of the enzyme is significantly lower than in matched individuals who have EOAD; indicating that trisomy of Hsa21 impairs enzyme activity during end-stage disease. These changes occur independently of neurofibrillary tangle stage, or the colocalisation of the enzyme with IBA1^+^ microglia and may result from the additional copy of cystatin B (*CSTB*) carried by individuals who have DS.

To investigate this further, we used mouse and human cellular preclinical models, that have an additional copy of *CSTB* which encodes cystatin B. In these model systems we observed that the abundance of cystatin B (CSTB) protein is increased but that cathepsin B enzyme activity is not robustly altered. We also observed this in the cortex of a DS-AD mouse model (*App*^NL-F/NL-F^; Dp(10)2Yey) at 3-months of age, which expresses human amyloid-β but has yet to accumulate significant aggregated peptide [8, 27]. Previous studies using aged transgenic *APP* mouse models have indicated an increase in cathepsin B protein abundance [32]. However, consistent with our results enhanced cathepsin B activity was also not observed in another amyloid-β accumulation model at 6-months of age [25]. In this study by Yang and colleagues, loss of a copy of *Cstb* was sufficient to increase cathepsin B activity in the mouse brain, but here we find conversely that an additional copy of *Cstb* is not sufficient to decrease cathepsin B activity. This is consistent with our previous report that duplication of the *Cstb* gene is also not sufficient to alter cathepsin B activity in the cortex of a mouse model, and underlies that loss and gain of function of a gene do not always have inverse consequences [28].

A previous study has reported a small but significant increase in cathepsin B activity in fibroblasts isolated from infants and children who had DS and typically developing controls [19]. We note that two of our trisomy 21 fibroblast lines (AG07438 and AG05397) and all of our disomic control lines differ from the lines studied by Jiang and colleagues, thus biological variation between the cells line used may contribute to the differences in our results. We note that our cellular preclinical model will not recapitulate cell-type specific functions of cystatin B (CSTB), such as its role at the synapse in neurons [33] or proposed regulatory role in microglia [34].

Our data indicate that the raised abundance of cystatin B (CSTB) protein is not sufficient to modulate cathepsin B activity in a range of model systems, perhaps because of the homeostatic regulation of the interaction of enzyme and inhibitor. However, our preclinical models do not recapitulate all aspects of end-stage (Braak VI) AD, notably the development of tau neurofibrillary pathology, a neuroinflammatory reaction to neuropathology and changes to the blood brain barrier, all of which may contribute to the dramatic increase in cathepsin B activity we observed in individuals who have EOAD. Thus, further research is required to determine if raised cystatin B (CSTB) may contribute to altered AD-associated cathepsin B activity in the context of these key aspects of the disease. This is important, as the changed AD-associated cathepsin B activity we observe, could contribute to differences in the progression of AD in people who have DS. Moreover, understanding why the activity is different in the presence of an additional copy of Hsa21 could provide significant novel insight into the underlying biology. Furthermore, our data indicate that targeting cathepsin B activity in people who have DS (i.e., an additional copy of Hsa21) is unlikely to have the same therapeutic effect as in individuals who have AD in the general population and that alternative targets should be considered for this important group of individuals who have a greatly elevated risk of early-onset disease.

Here we find a trend for cathepsin D activity to be decreased in the brains of individuals who have AD-DS compared to disomic healthy ageing controls. A previous study had shown that cathepsin D activity is robustly decreased in fibroblasts isolated from individuals who have DS compared to disomic controls [19]. Moreover, in a study of human post-mortem brain by Curtis and colleagues, cathepsin D activity was observed to be decreased in older adults who had DS (40-65 years of age), consistent with our observation [35]. They found that activity was not altered in younger adults who had DS (15-40 years of age) or in the brains of older adults (77-91 years of age) from the general population who had Late-onset AD (LOAD). Another report suggests a modest increase in cathepsin D activity occurs in younger individuals who have DS (2-45 years of age) [36], further studies are required to investigate these differences. Additionally, the number of cathepsin D positive cells, the abundance of cathepsin D transcript and protein is increased in LOAD [37–39]. Further research is required to understand how cathepsin D abundance and cellular location affects its function over the course of disease. Our data indicate that disomic individuals who have EOAD also have decreased cathepsin D activity, this suggests that decreased enzyme activity may be attributable to the early development of AD in mid-life rather than a direct effect of three copies of Hsa21 and these processes may differ in EOAD (caused by either an additional copy of Hsa21 or other mechanisms) and LOAD. This could relate to the elevated abundance of amyloid-β or APP-C-terminal fragment, which have been proposed to inhibit cathepsin D activity [19, 40] and also underlies the early development of AD-DS [3]. Additional studies to quantify enzyme activity across disease development, in independent cases, and correlate this with amyloid-β or APP-C-terminal fragment abundance are needed to verify this hypothesis.

## Study limitations

Here we study a case series of individuals who had EOAD and did, or did not, have DS, compared with healthily ageing individuals from the general population. As outlined, the demographics of the groups differed. In particular, the healthy ageing group was older than the AD-DS and EOAD groups, and the Braak tangle stage of the AD-DS and EOAD groups were not perfectly matched. To address these limitations, we included age at death as a covariant in our analysis and performed a sub-analysis of our Braak VI stage cases. In addition, here we only studied 10 individuals of each case-type and thus intra-individual variability may confound our results, and confirmation of our key findings in independent cases would be beneficial. Here we tested our hypothesis that raised cystatin B (CSTB) protein leads to reduced cathepsin B activity, as suggested by our human post-mortem analysis, in both human cellular and mouse models using a range of technical approaches. As outlined above, both preclinical systems have limitations and do not model all aspects of the biology of the human brain.

## Conclusion

Our data indicate both similarities and differences between EOAD in people who have and do not have DS. We find that cathepsin B activity is lower in people with DS when they develop AD compared with disomic individuals. In contrast, EOAD is associated with a similar decrease in cathepsin D activity in people with and without an additional copy of Hsa21. We also find that the number of cathepsin B^+^ cells and the abundance of cathepsin B protein does not differ between AD-DS and EOAD cases. However, the abundance of cystatin B (CSTB) is significantly higher in the temporal cortex of cases of AD-DS compared with EOAD or healthy ageing. We also find that in both a human cellular and mouse model of DS, cystatin B (CSTB) protein levels are increased but this is not sufficient to change cathepsin B activity. Research to understand these similarities and differences will both ensure that individuals who have DS have access to the most appropriate AD therapeutics and provides unique insight into the biology of AD pathogenesis by understanding how an additional copy of genes on Hsa21 can modulate disease.

## Abbreviations

AD: Alzheimer’s disease
AD-DS: Alzheimer’s disease in individuals who had Down syndrome
AFC: amino-4-trifluoromethyl coumarin
ALLM: N-Acetyl-L-leucyl-L-leucyl-L-methioninal
APP: amyloid precursor protein
CatB: cathepsin B
CatD: cathepsin D
CSTB: cystatin B
DMEM: Dulbecco’s Modified Eagle Medium
DS: Down syndrome
EOAD: early-onset Alzheimer’s disease
EPM1: Unverricht-Lundborg type
MCA: 7-Methoxycoumarin-4-acetic acid
PMI: post-mortem interval
SEM: standard error of the mean
WT: wildtype

## Declarations

### Ethical approval and consent to participate

#### Human tissue

The procurement and use of human tissues in this study was in accordance with the UK Human Tissue Act 2004. The study was reviewed and approved by NHS Research Ethics committee, London-Queen Square (REC 09/H0716/57). All samples were supplied, anonymized by the South West Dementia Brain Bank, Bristol University, UK, and had full research consent (REC 18/SW/0029).

#### Animal research

All experiments were undertaken in accordance with the Animals (Scientific Procedures) Act 1986 (United Kingdom), after local institutional ethical review by the Medical Research Council, University College London under licence from the UK government and reported in accordance with ARRIVE 2.0 guidelines.

#### Consent to publish

Not applicable

## Availability of data and material

The datasets generated during the current study available from the corresponding author on reasonable request.

## Author Contributions

E.M.C.F., and F.K.W. designed the research and applied for ethical permission for research; Y.W., A.M., D.L., P.M., S.N., K.C., and F.K.W. performed research; Y.W., K.C., P.M., and F.K.W. analysed data; Y.W., P.M., D.L. S.N., K.C., F.D., M.V., A.M., E.M.C.F., and F.K.W. edited the paper; F.K.W. wrote the paper; F.D. and M.V. contributed reagents/analytic tools and procedures.

## Acknowledgments

The authors thank Eugene Yu (Rosewell Cancer Research Institute, USA) for Dp(10)2Yey and Dp(17)3Yey mouse models and Takaomi Saido (Riken Institute, Japan) for the *App^NL-F/NL-F^* mouse model. We thank the Coriell Cell repositories, USA for the human fibroblast cell lines used in this study. We would like to thank the South West Dementia Brain Bank (SWDBB), and the generosity of the donors and their families for providing brain tissue for this study. We would also like to thank Connor Scott (UCL) for offering valuable guidance regarding the process of immunostaining. We thank Oke Avwenagha (UCL) for her assistance with this project. For the purpose of Open Access, the authors have applied a CC BY public copyright licence to any Author Accepted Manuscript version arising from this submission.

## Supplementary Data

**Supplementary figure 1.**
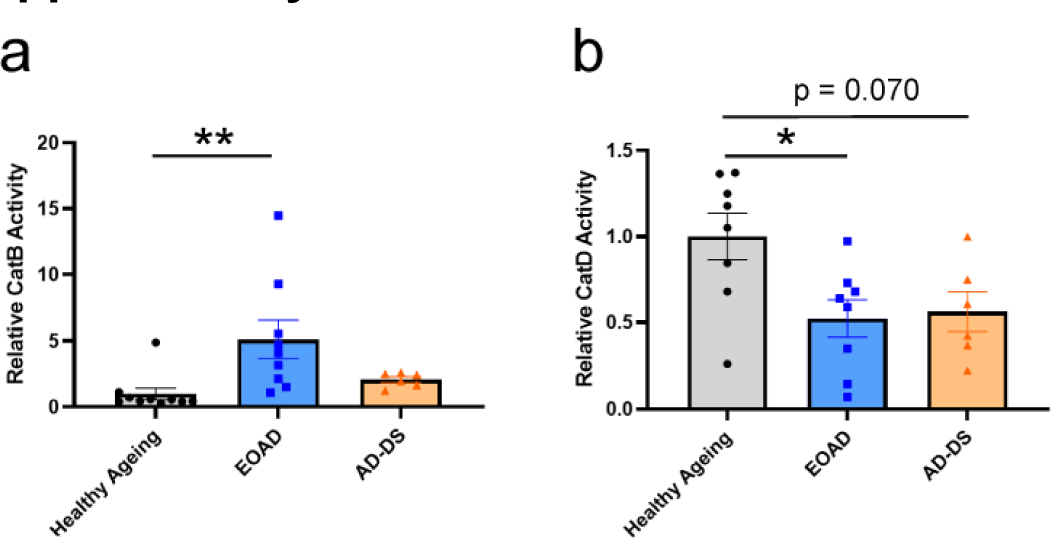
Changes to cathepsin B and cathepsin D activity in a sub analysis of EOAD and AD-DS Braak VI cases. Sub-analysis of cathepsin B (a) and (b) cathepsin D activity in EOAD and AD-DS (Braak neurofibrillary tangle stage VI), and healthy-ageing. a) Type of case affected cathepsin B activity (ANOVA F(2,17) = 6.792, p = 0.007); activity was significantly higher in individuals who had EOAD than in controls (pairwise comparisons with Bonferroni correction p = 0.008), with no difference between individuals with AD-DS and control individuals (pairwise comparisons with Bonferroni correction p = 1.000) or AD-DS and EOAD (pairwise comparisons with Bonferroni correction p = 0.115). There was a trend that the age at the time of death (in years) has an impact on activity (ANOVA F (1,17) = 4.358, p = 0.053) (Healthy ageing n = 9, EOAD n = 9, AD-DS n = 6). (**b**) Type of case (ANOVA F(2,14) = 7.188, p = 0.007) and age at death (ANOVA F(2,14) = 5.214, p = 0.039) significantly affected cathepsin D activity. Activity was significantly lower in individuals who had EOAD than in healthy controls (pairwise comparisons with Bonferroni correction p = 0.028), a trend for reduced activity was observed between AD-DS and healthy ageing (pairwise comparisons with Bonferroni correction, p = 0.070), with no difference in the activity between individuals with AD-DS and those with EOAD (pairwise comparisons p = 1.000)) (Healthy ageing n = 8, EOAD n = 8, AD-DS n = 6). Individual data points are technical means for independent biological samples, error bars SEM. * p < 0.05 and ** p < 0.01.

**Supplementary table 1.** Samples of human post-mortem temporal cortex (BA21) used for this research, including UK Brain Bank Identification Number (UK BBN ID), and anonymised demographics of the cases studied including Braak neurofibrillary tangle stage and *APOE* genotype, as reported by the SWDBB brain bank.

| UK BBN ID | Gender | Age at death (years) | PMI (hours) | APOE | Status | Braak |
| --- | --- | --- | --- | --- | --- | --- |
| BBN_9061 | F | 54 | 24 | 3,3 | AD | VI |
| BBN_9162 | M | 63 | 43 | 3,3 | AD | VI |
| BBN_9212 | F | 65 | 22 | 3,3 | AD | VI |
| BBN_9259 | F | 47 | 54 | 3,4 | AD | VI |
| BBN_9265 | F | 60 | 5 | 2,3 | AD | VI |
| BBN_9291 | M | 62 | 25 | 3,4 | AD | VI |
| BBN_9315 | F | 67 | 24 | 4,4 | AD | VI |
| BBN_9342 | F | 65 | 12 | 4,4 | AD | VI |
| BBN_4202 | M | 64 | 67 | 4,4 | AD | V |
| BBN_9406 | M | 61 | 33 | 2,3 | AD | VI |
| BBN_8761 | M | 63 | 31 | 3,3 | DS-AD | n/a |
| BBN_8867 | M | 62 | 24 | 3,4 | DS-AD | VI |
| BBN_8869 | M | 62 | 51 | 3,3 | DS-AD | V |
| BBN_8881 | F | 63 | 51 | 2,3 | DS-AD | VI |
| BBN_8887 | F | 50 | 43 | 3,4 | DS-AD | VI |
| BBN_8953 | F | 64 | 48 | 3,3 | DS-AD | VI |
| BBN_8991 | M | 64 | 16 | 3,3 | DS-AD | V |
| BBN_8994 | F | 59 | 24 | 3,3 | DS-AD | VI |
| BBN_9097 | F | 67 | 17 | 3,4 | DS-AD | VI |
| BBN_9184 | F | 48 | 79 | 2,4 | DS-AD | IV |
| BBN_8696 | M | 63 | 40 | 3,3 | Healthy-ageing |  |
| BBN_8700 | M | 64 | 12 | 3,4 | Healthy-ageing |  |
| BBN_8702 | M | 58 | 20 | 2,3 | Healthy-ageing |  |
| BBN_8703 | M | 64 | 16 | 3,3 | Healthy-ageing |  |
| BBN_8835 | F | 73 | 59 | 3,3 | Healthy-ageing |  |
| BBN_8980 | F | 72 | 24 | 3,3 | Healthy-ageing |  |
| BBN_9389 | F | 68 | 39 | 2,3 | Healthy-ageing | 0 |
| BBN_9399 | F | 73 | 50 | 3,4 | Healthy-ageing | II |
| BBN_9422 | F | 74 | 40 | 3,3 | Healthy-ageing | I |
| BBN_9432 | F | 74 | 28 | N/A | Healthy-ageing | II |

**Supplementary table 2.**
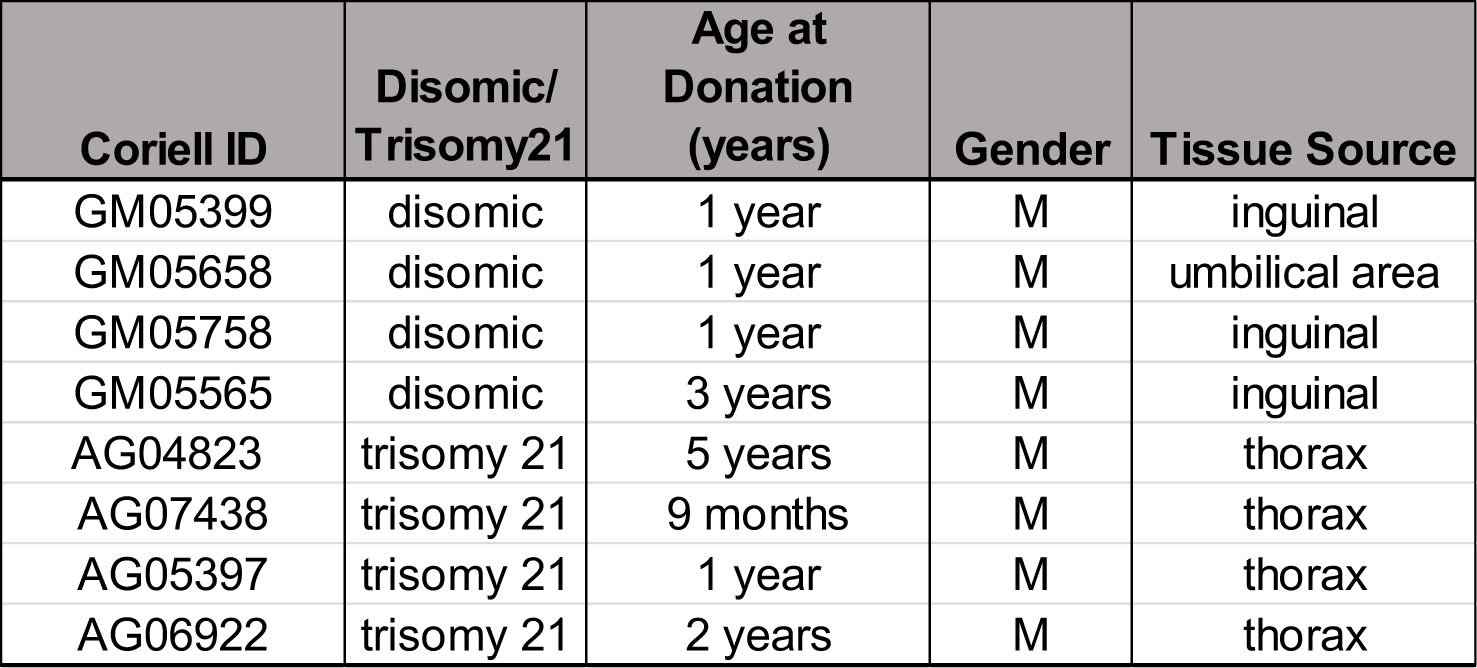
Demographics for the human fibroblast lines used in this research, including Coriell Institute for Medical Research ID, and anonymised details of the donor.

**Supplementary table 3.** Summary information of the number and sex of individual mice used in this research.

| Cohort | WT (n) |  | Segmental<br>duplication (n) |  | *App*<sup>NL-F/NL-F</sup> (n) |  | Segmental<br>duplication; *App*<sup>NL-F/NL-F</sup><br>(n) |  |
| --- | --- | --- | --- | --- | --- | --- | --- | --- |
|  | male | female | male | female | male | female | male | female |
| Dp(10)2Yey * *App*<sup>NL-F/NL-F</sup> | 4 | 1 | 8 | 0 | 9 | 0 | 4 | 2 |
| Dp(10)2Yey (ALLM) | 3 | 5 | 4 | 2 | N/A | N/A | N/A | N/A |
| Dp(10)2Yey (FMK) | 3 | 3 | 8 | 5 | N/A | N/A | N/A | N/A |
| Dp1Tyb | 8 | 6 | 6 | 8 | N/A | N/A | N/A | N/A |
| Dp(17)3Yey | 4 | 4 | 4 | 4 | N/A | N/A | N/A | N/A |

